# Toll-like receptor expression across histological states of Epidermodysplasia Verruciformis

**DOI:** 10.64898/2026.09.29.755296

**Authors:** Luis Alberto Ribeiro Fróes Júnior, Cibele Conceição dos Apóstolos Pereira, Lana Luiza da Cruz Silva, Maíra Garcia Saldanha, Naiura Vieira Pereira, Walmar Roncalli Pereira de Oliveira, Mirian Nacagami Sotto

## Abstract

Epidermodysplasia verruciformis (EV) is characterized by susceptibility to cutaneous β-human papillomaviruses, persistent viral lesions and an increased risk of cutaneous squamous cell carcinoma (cSCC). We investigated whether expression of Toll-like receptors 3, 4, 5 and 9 (TLR3, TLR4, TLR5 and TLR9) differs across histological states within EV. Immunoreactivity was quantified by digital H-score using QuPath in 97 specimens from 28 patients, comprising 13 lesion-free skin samples, 15 flat warts and 69 cSCCs. Prespecified comparisons evaluated flat wart versus lesion-free skin, cSCC versus non-neoplastic tissue and invasive versus in situ cSCC, with invasive tumour differentiation assessed secondarily. Patient-clustered generalized estimating equations and false-discovery-rate (FDR) correction were used for inference. After correction for multiple testing, TLR9 H-scores were higher in flat warts than in lesion-free skin (β=60.58; q=0.0332). Compared with non-neoplastic tissue, cSCC showed lower TLR3 (β=−81.40; q=0.00745), higher TLR4 (β=55.59; q=0.04793) and lower TLR9 expression (β=−34.80; q=0.03412). TLR5 showed a nominal increase in cSCC that did not meet the FDR threshold. None of the four receptors differed significantly between invasive and in situ cSCC, and no differentiation trend remained significant after FDR correction. These findings show receptor-specific differences in TLR expression across histological states of EV, particularly involving TLR3, TLR4 and TLR9, although the diagnostic and prognostic significance of these tissue patterns remains to be established.

## Introduction

Epidermodysplasia verruciformis (EV) is a rare disorder characterized by selective susceptibility to cutaneous β-human papillomaviruses (β-HPVs), persistent flat warts and a marked predisposition to cutaneous squamous cell carcinoma (cSCC), particularly on sun-exposed skin (Béziat et al. 2021; Smola 2020). In contrast to the sustained oncogenic role of high-risk mucosal HPV, β-HPVs are thought to cooperate with ultraviolet radiation and host susceptibility, particularly during earlier stages of cutaneous carcinogenesis (Smola 2020). EV therefore provides a distinctive human context linking persistent β-HPV infection, epithelial host responses and keratinocyte neoplasia.

Keratinocytes actively participate in cutaneous innate immunity and functionally express TLR3, TLR4, TLR5 and TLR9 (Lebre et al. 2007). These receptors sense distinct classes of innate immune ligands: TLR3 and TLR9 recognize double-stranded RNA and unmethylated CpG-rich DNA, respectively, whereas TLR4 and TLR5 classically respond to lipopolysaccharide and flagellin (Lebre et al. 2007). Activation of these receptors in human keratinocytes induces distinct but partially overlapping cytokine and chemokine responses, with TLR3 and TLR9 ligands additionally inducing type I interferons (Lebre et al. 2007). Beyond pathogen sensing, TLR4 expression has been shown experimentally to increase with keratinocyte differentiation, while experimental manipulation of TLR4 altered keratinocyte proliferation (Iotzova-Weiss et al. 2017).

Human cutaneous lesions likewise show context-dependent TLR profiles. Ku et al. (2008) reported increased TLR3 and TLR9 mRNA expression and epidermal immunostaining in verruca vulgaris and molluscum contagiosum relative to normal skin. In cSCC, TLR3 and TLR5 transcripts were higher in tumour centres than in distant normal skin (Muehleisen et al. 2012). TLR4 immunoreactivity was reported to be higher in cSCC than in actinic keratosis or Bowen disease and lower in poorly differentiated than in moderately or well-differentiated tumours (Mikami et al. 2019). More recent work has also shown heterogeneous TLR3 protein staining in cSCC, without a consistent relationship to histological subtype (Schneider et al. 2024). These observations support examining individual TLRs across viral and neoplastic cutaneous contexts rather than assuming a uniform pattern across receptors.

EV-specific evidence remains limited. Fróes et al. (2025) reported lower TLR3 and TLR9 immunoreactivity in both lesion-free skin and flat warts from patients with EV than in corresponding tissues from individuals without EV. In cSCC, Pereira et al. (2026) found higher TLR4 and TLR9 immunoreactivity in EV-associated than in non-EV tumours. Together, these studies do not provide a single analytical framework examining TLR3, TLR4, TLR5 and TLR9 across lesion-free skin, flat warts, cSCC in situ and invasive cSCC within an EV cohort. We therefore investigated whether immunoreactivity for these four receptors differed across these histological states and, secondarily, whether receptor immunoreactivity was associated with tumour differentiation among invasive cSCCs.

## Materials and Methods

### Study design, specimens and ethics

This retrospective study analysed archived skin specimens from patients with epidermodysplasia verruciformis (EV). Specimens were classified as lesion-free skin, flat wart, cSCC in situ, or invasive cSCC of histological grade 1, 2 or 3. Both conventional whole sections and tissue microarrays (TMAs) were analysed, with no individual carcinoma represented in both formats. Distinct specimens from the same patient were retained as separate observations. For tumours represented by multiple TMA cores, core-level measurements were aggregated at the tumour level as described below. Some patients in this cohort had undergone β-HPV typing in previous studies of the Brazilian EV series, but comprehensive lesion-level genotype data were not available for the specimens included in the present analysis. The study was approved by the Institutional Review Board of Hospital das Clínicas, University of São Paulo Medical School (protocol 50871415.2.0000.0068; approval no. 1.401.205, 3 February 2016), in accordance with the Declaration of Helsinki. The requirement for informed consent was waived because the study exclusively used archived, anonymized specimens.

### Immunohistochemistry

Primary antibodies were TLR3 (IMGENEX, San Diego, CA, USA; IMG-315A; clone 40C1285.6; 1:100), TLR4 (eBioscience, San Diego, CA, USA; clone HTA125; 1:50), TLR5 (IMGENEX; IMG-664A; clone 19D759.2; 1:200) and TLR9 (IMGENEX; IMG-305A; clone 26C593.2; 1:200). Sections 3–4 μm thick were deparaffinized and rehydrated, followed by antigen retrieval in citrate buffer (pH 6.0) at 95°C for 20 min in a pressure cooker. Endogenous peroxidase activity was blocked with 3% hydrogen peroxide in methanol for 10 min. Detection was performed using the Novolink Max Polymer Detection System (Leica Microsystems, Newcastle, UK; RE7280-K), with 3,31l-diaminobenzidine (DAB; Sigma Chemical Company, St. Louis, MO, USA; D5637) as chromogen. Sections were counterstained with Harris haematoxylin for 10 s, dehydrated and mounted with Permount (Fisher Scientific, Fair Lawn, NJ, USA; SP15-100).

Positive controls comprised normal skin and molluscum contagiosum for TLR3, TLR4 and TLR5, and cutaneous leishmaniasis for TLR9. For negative controls, primary antibodies were replaced with isotype-matched non-immune serum. Current manufacturer documentation for the corresponding clones lists immunohistochemistry among their applications: TLR3 clone 40C1285.6 (Novus/Bio-Techne NBP2-24875), TLR4 clone HTA125 (Thermo Fisher/eBioscience 14-9917-82; RRID AB_468607), TLR5 clone 19D759.2 (Novus/Bio-Techne NBP2-24787) and TLR9 clone 26C593.2 (Novus/Bio-Techne NBP2-24729).

### Digital slide acquisition and region selection

The digital image set comprised 51 immunohistochemistry whole-slide images from conventional sections and eight TMA images corresponding to two TMA blocks (T0351 and T0352) stained for TLR3, TLR4, TLR5 and TLR9. All quantitative immunohistochemistry slides were digitized using the same Aperio ScanScope AT scanner and stored in SVS format. For the 51 conventional-section images, project metadata recorded a nominal magnification of 40×, an x–y pixel calibration of 0.2476 μm/pixel and RGB representation at 8 bits per channel. Separate TMA haematoxylin-and-eosin images acquired on a 3DHistech system were used for morphological orientation only and were not included in the quantitative analysis.

Each available conventional-section sample–TLR combination was represented by one ROI, whereas eligible annotated TMA cores were quantified separately. Areas of tissue loss, folds, necrosis or other non-analysable material were excluded. ROI delineation and review were completed before H-score generation and without reference to the assigned analytical histological category.

### Digital image analysis and H-score

Digital image analysis was performed in QuPath version 0.7.0. Images were defined as brightfield haematoxylin–DAB, and the default QuPath H-DAB colour-deconvolution settings were applied unchanged to all markers and image formats. Nuclei were detected within eligible annotations using Positive Cell Detection with Haematoxylin OD as the detection image to define individual cells. Marker-specific nuclear-detection parameters were a threshold of 0.050 and background radius of 8 μm for TLR3, 0.050 and 0 μm for TLR4, 0.050 and 0 μm for TLR5, and 0.075 and 8 μm for TLR9. All markers otherwise used a requested pixel size of 0.5 μm, background reconstruction, median radius of 0 μm, sigma of 1.5 μm, nuclear area of 10–400 μm^2^, maximum background of 2, watershed post-processing, exclusion of DAB from nuclear detection, cell expansion of 5 μm, inclusion of nuclei and boundary smoothing.

Per-cell DAB intensity was measured as *Cytoplasm: DAB OD mean*. Intensity thresholds were calibrated separately for each TLR using visually selected 0/1+/2+/3+ reference regions and exported single-cell DAB optical-density distributions. For TLR3, TLR4 and TLR5, the negative/1+ threshold was anchored approximately to the 95th percentile of the visually negative reference, with the positive-category thresholds guided by the distributions of adjacent reference classes. For TLR9, because the 1+ and 2+ reference distributions overlapped, the 1+ anchor was derived from REF1 values above the negative threshold, whereas the 2+ anchor was based on the upper quartile of REF2; the resulting cutoffs were subsequently validated visually in mixed-intensity regions. Calibration was completed before analytical H-score generation and without reference to the assigned analytical histological category.

The final thresholds separating negative/1+, 1+/2+ and 2+/3+ cells were 0.099/0.129/0.175 for TLR3, 0.123/0.184/0.230 for TLR4, 0.076/0.151/0.231 for TLR5 and 0.146/0.171/0.211 for TLR9. For each marker, these thresholds were subsequently applied without adjustment to the other TMA block and to conventional sections, with visual confirmation of correspondence of the 0/1+/2+/3+ intensity classes before batch reprocessing. No colour-normalization procedure was applied.

For each ROI, H-score was calculated as 1 × (% 1+) + 2 × (% 2+) + 3 × (% 3+), yielding a score from 0 to 300; all detected cells classified as negative, 1+, 2+ or 3+ contributed to the denominator. For each TMA tumour, eligible core-level H-scores were summarized by their median, yielding one H-score per tumour and TLR. Conventional-section samples were represented by the H-score of their single ROI.

### Quality control and data integrity

Final segmentation settings were established after visual review for missed, fragmented or fused nuclei, artefactual detections and adequacy of cell expansion. A prespecified minimum of 100 detected cells per ROI or TMA core was required for the primary analysis. Two TMA cores fell below this threshold and were excluded when deriving the primary tumour-level H-scores but reintroduced in a prespecified sensitivity analysis.

The final export was reconciled against the image/ROI inventory. For each ROI, negative, 1+, 2+ and 3+ cell counts summed to the total number of detected cells, and H-scores were independently recalculated from the exported intensity-class data. Sample identities were reconciled against the original source records, and TMA core positions and tumour-level groupings were cross-checked against the physical TMA maps and clinical tumour-selection records

### Statistical analysis

TLR3, TLR4, TLR5 and TLR9 were analysed separately using continuous H-score as the outcome. For each marker, three primary contrasts were prespecified and evaluated in separate models: (1) flat wart versus lesion-free skin; (2) cSCC (in situ and invasive grades 1–3) versus non-neoplastic tissue (lesion-free skin and flat wart); and (3) invasive cSCC (grades 1–3) versus cSCC in situ. Invasive tumour differentiation was evaluated secondarily using a 1-degree-of-freedom linear trend with grades coded G1=1, G2=2 and G3=3. To assess whether inference depended on the linearity assumption, grade was also modelled categorically in a global G1/G2/G3 test. These comparisons were interpreted as cross-sectional differences between histological states or grades, not as within-patient longitudinal transitions.

Marginal models were fitted using Gaussian generalized estimating equations with identity link, patient as the clustering variable and working independence as the primary correlation structure. Regression coefficients were expressed in H-score points. For the three primary binary contrasts, β represents the marginal mean difference between groups; for the linear differentiation trend, β represents the mean change in H-score per one-grade increment. Primary inference used cluster-robust CR2 variance estimation, with Satterthwaite small-sample degrees of freedom for single-parameter tests and HTZ tests for multiparameter hypotheses. Working exchangeable correlation with the same CR2-based inference was assessed as a sensitivity analysis. A separate small-sample sensitivity analysis used the Fay– Graubard d5 adjustment with bound 0.75. Analyses that were mathematically non-estimable were reported as such without post hoc substitution of an alternative inferential method.

Primary models were unadjusted. A prespecified covariate-adjusted sensitivity analysis included age as a continuous variable and photoexposure. Following review of covariate provenance, this analysis was implemented as a complete-case analysis restricted to directly documented photoexposure; an additional sensitivity analysis also allowed proxy-derived photoexposure values. Missing H-scores were not imputed, and structurally unavailable marker measurements were not treated as zero.

Multiplicity in the primary inferential family was controlled using the Benjamini–Hochberg false-discovery rate (FDR) at q=0.05 across 12 tests (three primary contrasts × four TLRs). The four linear differentiation-trend tests constituted a separate FDR family, and the four categorical global grade tests constituted a third family. Prespecified TMA sensitivity analyses reintroduced cores below the 100-cell threshold, replaced the median core-level summary with either the arithmetic mean or the cell-count-weighted mean, and separately restricted TMA analyses to tumours represented by at least two eligible cores. Analyses were performed using geepack 1.3.13, clubSandwich 0.7.0 and saws 0.9.7.0.

### Use of generative AI

During preparation of the methodological record and manuscript, the authors used ChatGPT (OpenAI) to assist with the organization of methodological information, drafting of analysis code, and drafting and revision of manuscript text. All AI-assisted outputs were reviewed and verified by the authors against the underlying sources and data. The authors reviewed the final analysis code, statistical results and manuscript text and take full responsibility for the content of the manuscript.

## Results and Discussion

### Analytical cohort and marker availability

The analytical EV cohort comprised 97 specimens from 28 patients: 13 lesion-free skin specimens, 15 flat warts and 69 cutaneous squamous cell carcinomas (cSCCs). The cohort included 16 women and 12 men; age at biopsy was available for 96 specimens and ranged from 8 to 70 years. Fifty-eight specimens were classified as photoexposed and 38 as non-photoexposed, with photoexposure unavailable for one flat wart. None of the patients was known to have HIV infection or another systemic immunodeficiency. Nineteen patients contributed carcinomas; 47 cSCCs were represented by TMA cores and 22 by conventional whole sections. After tumour-level aggregation of TMA core measurements, marker-specific H-scores were available for 93 specimens for TLR3, 69 for TLR4, 87 for TLR5 and 95 for TLR9. Representative immunohistochemical staining patterns of TLR3, TLR4, TLR5 and TLR9 in lesion-free skin, flat warts, cSCC in situ and invasive cSCC are shown in Fig. 1.

**Fig 1.**
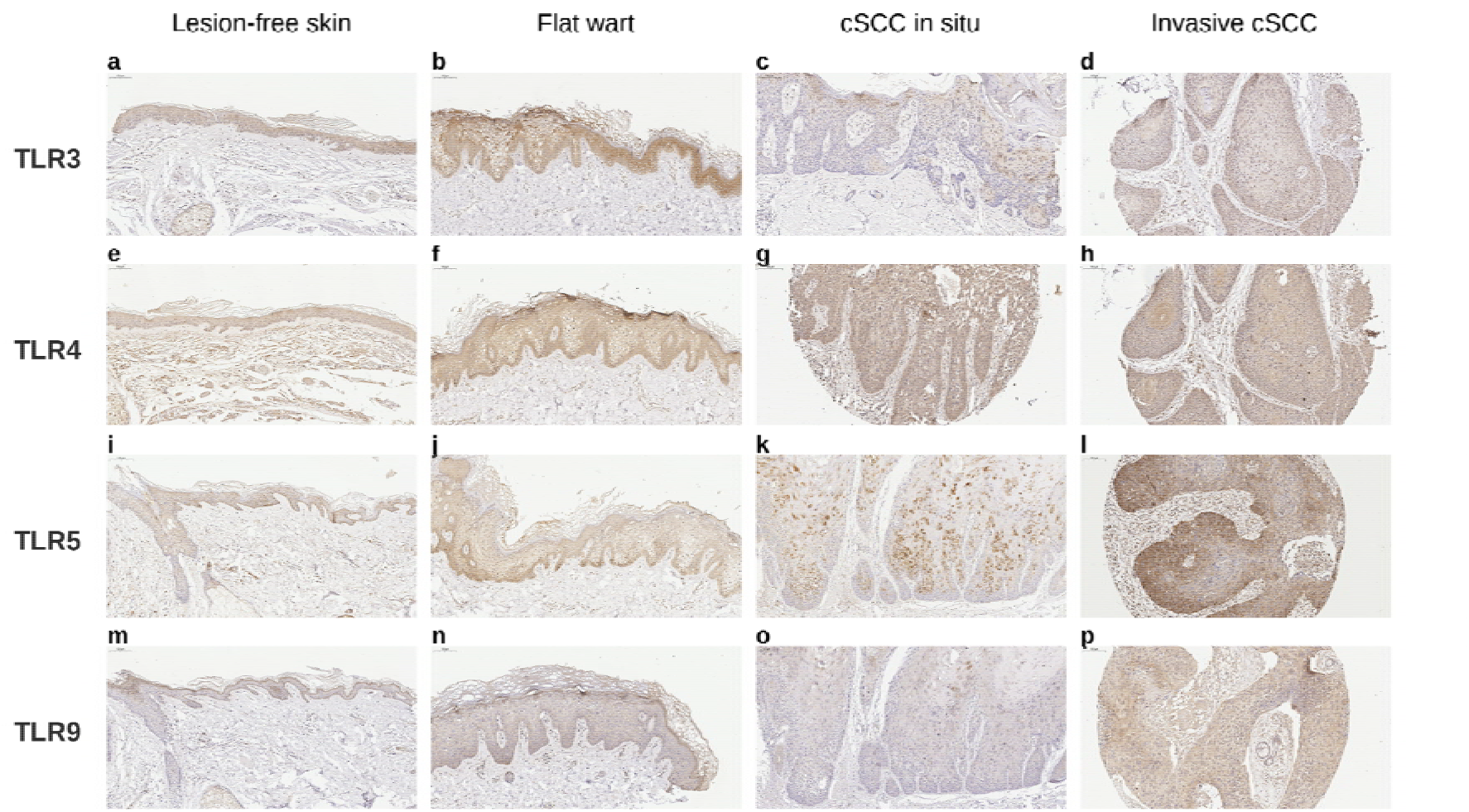
Representative immunohistochemical staining patterns of TLR3, TLR4, TLR5 and TLR9 in epidermodysplasia verruciformis skin specimens. a–d TLR3, e–h TLR4, i–l TLR5 and m–p TLR9. Within each row, panels show lesion-free skin, flat wart, cSCC in situ and invasive cSCC, respectively. Haematoxylin counterstain; scale bars, 100 μm.

### TLR expression across histological states

H-score distributions across lesion-free skin, flat warts and cSCC are shown in Fig. 2. Four of the 12 prespecified primary contrasts remained significant after BH-FDR correction (Fig. 3). Compared with non-neoplastic EV tissue (lesion-free skin plus flat wart), cSCC showed lower TLR3 H-scores (β = −81.40, 95% CI −117.24 to −45.56, q=0.0075), higher TLR4 H-scores (β = 55.59, 95% CI 13.77 to 97.41, q=0.0479) and lower TLR9 H-scores (β = −34.80, 95% CI −58.66 to −10.95, q=0.0341). Flat warts also showed higher TLR9 H-scores than lesion-free skin (β = 60.58, 95% CI 22.33 to 98.84, q=0.0332).

**Fig 2.**
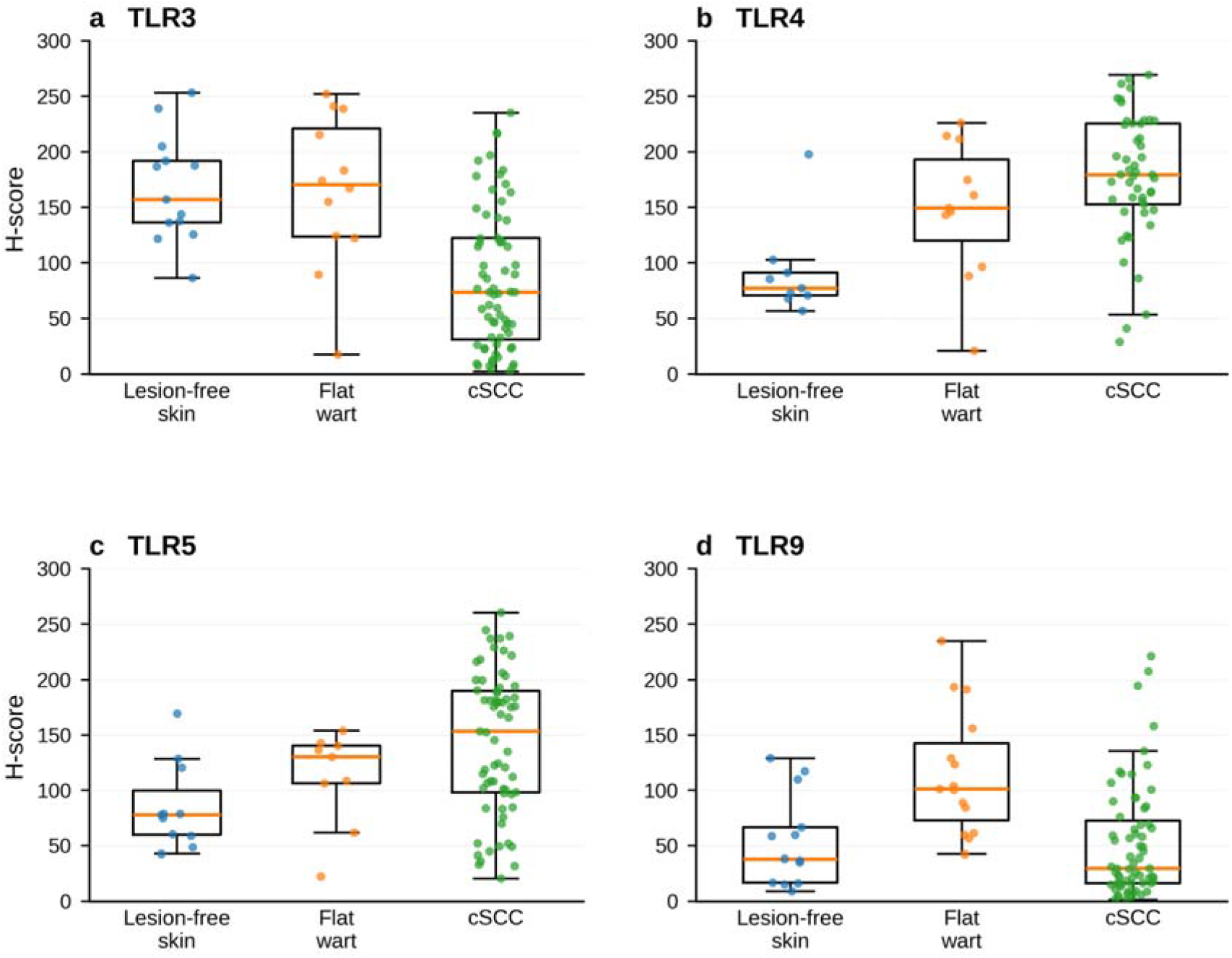
Distribution of TLR3 (a), TLR4 (b), TLR5 (c) and TLR9 (d) H-scores in lesion-free skin, flat warts and cutaneous squamous cell carcinoma (cSCC). cSCC comprises in situ and invasive grade 1–3 tumours. Each point represents one analytical specimen H-score; for TMA tumours represented by multiple eligible cores, the plotted value is the median core H-score. Boxes indicate the interquartile range, central lines the median and whiskers the most extreme values within 1.5 times the interquartile range.

**Fig 3.**
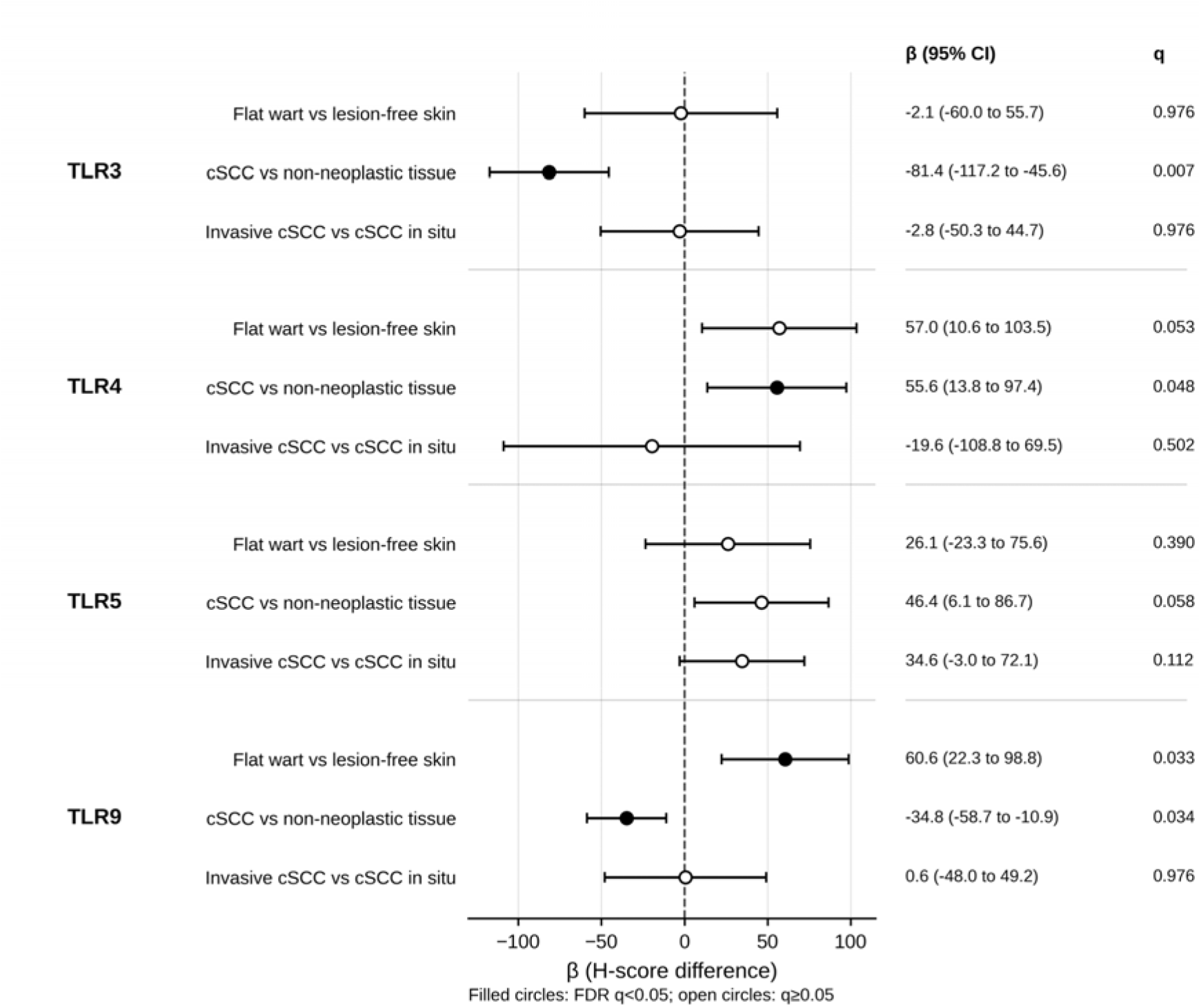
Estimated H-score differences for the 12 prespecified primary contrasts. Points indicate marginal mean differences (β) from Gaussian generalized estimating equations, with 95% confidence intervals based on CR2 cluster-robust variance estimation with Satterthwaite small-sample degrees of freedom. Positive β values indicate higher H-scores in the first-named group and negative values lower H-scores. Filled circles denote contrasts significant after Benjamini–Hochberg false-discovery-rate correction across the 12-test primary family (q<0.05); open circles denote q≥0.05. The dashed vertical line indicates β=0 The three FDR-significant cSCC-related associations retained their direction, broadly similar effect estimates and nominal significance across the prespecified TMA-core inclusion and aggregation sensitivity analyses; the TLR9 flat-wart-versus-lesion-free-skin contrast did not involve TMA-derived observations and was therefore unchanged in these analyses. The four FDR-significant primary associations also retained their direction and nominal significance under an exchangeable working correlation and Fay–Graubard d5 inference.

TLR4 H-scores were higher in flat warts than in lesion-free skin (β = 57.04, p=0.0222, q=0.0533), and TLR5 H-scores were higher in cSCC than in non-neoplastic tissue (β = 46.42, p=0.0292, q=0.0584), but neither contrast remained significant after FDR correction. None of the invasive cSCC versus cSCC in situ contrasts was FDR-significant.

Covariate adjustment preserved the direction of all four primary associations. The TLR3 cSCC association and both TLR9 associations remained nominally significant in analyses using either directly documented photoexposure or documented plus proxy-derived values. For TLR4, the cSCC-versus-non-neoplastic estimate remained positive but was less precisely estimated after adjustment, with confidence intervals including zero both with directly documented photoexposure (β=41.80, 95% CI −12.78 to 96.38, p=0.113) and when proxy-derived photoexposure was also allowed (β=45.55, 95% CI −6.81 to 97.91, p=0.0791). Complete sensitivity-analysis results are provided in Online Resource 2.

The lower TLR3 H-scores in cSCC correspond to lower TLR3 immunoreactivity than in non-neoplastic EV tissue. Published TLR3 findings in cutaneous cSCC are not uniform. Muehleisen et al. (2012) reported increased TLR3 mRNA in cSCC tumour centres relative to distant normal skin, whereas Schneider et al. (2024) described heterogeneous TLR3 protein staining in cSCC without a consistent relationship to histological subtype. Experimental data for HPV8 do not support a simple model of reduced TLR3 abundance: HPV8 early genes had little overall effect on TLR3 expression, with HPV8 E2 associated with increased TLR3, despite HPV8 E1/E2 suppressing MDA5-induced type I interferon responses (Rattay et al. 2022). Direct comparison is limited by differences in analyte, quantification method and comparison groups; bulk tumour transcript abundance, semiquantitative immunohistochemistry and digital H-score are not directly equivalent measures.

The higher TLR4 H-scores in cSCC correspond to greater TLR4 immunoreactivity than in non-neoplastic EV tissue. This direction is broadly consistent with findings from other cutaneous cSCC settings. Janda et al. (2016) reported increased TLR4 staining in actinic keratoses and cSCC relative to normal skin, while Mikami et al. (2019) found higher TLR4 expression in cSCC than in actinic keratosis or Bowen disease. In a separate between-cohort comparison, Pereira et al. (2026) reported higher TLR4 expression in EV-associated than in non-EV cSCC. That comparison addresses a different question from the present within-EV contrast between cSCC and non-neoplastic tissue.

TLR9 immunoreactivity was higher in flat warts than in lesion-free skin but lower in cSCC than in the combined non-neoplastic group. Outside EV, increased epidermal TLR9 immunoreactivity and TLR9 transcript expression have been reported in verruca vulgaris and molluscum contagiosum relative to normal skin (Ku et al. 2008). More direct type-specific evidence is available for HPV38. HPV38 E7 promotes ΔNp73α-dependent repression of TLR9 transcription, while HPV38 E6/E7 inhibit UV-induced TLR9 upregulation by preventing recruitment of p53 and c-Jun to the TLR9 promoter (Pacini et al. 2015, 2017). These findings demonstrate direct HPV38-mediated interference with TLR9 regulation in experimental keratinocytes, but do not establish that HPV38 or any other individual β-HPV type accounts for the tissue-level TLR9 differences observed in the present cohort.

In a previous comparative EV study, TLR9 immunoreactivity was lower in EV than in non-EV skin, but no significant flat-wart-versus-lesion-free-skin difference was detected within EV (Fróes et al. 2025). That analysis quantified the percentage of DAB-positive epithelial area, whereas the present study used an intensity-weighted cellular H-score and patient-clustered inference. The two analyses therefore use different quantitative endpoints and inferential approaches. Pereira et al. (2026) also reported higher TLR9 expression in EV-associated than in non-EV cSCC; that between-cohort tumour comparison addresses a different question from the present comparison of cSCC with non-neoplastic tissue within EV.

The positive estimate for the TLR5 cSCC contrast is consistent in direction with Muehleisen et al. (2012), who reported higher TLR5 mRNA in cSCC tumour centres than in distant normal skin. However, the present contrast did not meet the FDR threshold, and the two studies assessed different analytes and comparison groups.

Across receptors, the differing patterns show that the relationship between TLR immunoreactivity and histological state is receptor-specific rather than uniform across EV tissues. TLR9 showed significant associations in opposite directions in the flat-wart-versus-lesion-free-skin and cSCC-versus-non-neoplastic comparisons. These cross-sectional tissue-level differences do not establish altered receptor signalling, causality or a temporal sequence of lesion evolution.

### Tumour differentiation

No linear G1⍰G2⍰G3 differentiation trend remained significant after FDR correction. TLR5 showed a positive linear trend at the nominal level, corresponding to 23.15 H-score points per grade increment (95% CI 0.48 to 45.81, p=0.0470), but this did not remain significant after correction of the four-test family (q=0.161). The categorical global G1/G2/G3 tests were also non-significant after their separate FDR correction for all four receptors. H-score distributions across lesion-free skin, flat wart, cSCC in situ and invasive grades G1–G3 are shown in Online Resource 1.

The negative TLR4 estimate across G1–G3 was directionally consistent with previous observations by Mikami et al. (2019), who reported lower TLR4 expression in poorly differentiated cSCC than in moderately or well-differentiated tumours. However, the present estimate was not statistically significant and therefore does not provide confirmatory evidence of a grade-related association. Experimentally, TLR4 expression has also been shown to increase with keratinocyte differentiation in vitro (Iotzova-Weiss et al. 2017), although experimental keratinocyte differentiation is not equivalent to histological differentiation of cSCC.

### Strengths and limitations

Several limitations should be considered. This retrospective study used archived material and had unequal marker-specific analytical sample sizes, with limited numbers in some histological strata. The cSCC dataset included both TMA-derived and conventional whole-section measurements, with no carcinoma represented in both formats; direct within-tumour assessment of concordance between TMA and whole-section H-scores was therefore not possible. Although the same marker-specific intensity thresholds were applied across both formats, differences in tissue sampling between TMA cores and whole sections cannot be excluded as a source of variability. β-HPV genotype data were not available with reliable specimen-level linkage across the analytical cohort, precluding genotype-stratified analysis of TLR immunoreactivity. Finally, H-score quantifies immunoreactivity rather than receptor signalling activity, and no functional assays were performed.

Strengths include acquisition of all quantitative IHC images on the same scanner, ROI delineation and review before H-score generation and without reference to the assigned analytical histological category, marker-specific intensity calibration before batch analysis, tumour-level aggregation of multiple eligible TMA cores, reconciliation of analytical data with source records, patient-clustered inference with multiplicity control, and sensitivity analyses addressing TMA aggregation, working correlation, small-sample inference and covariate adjustment.

TLR immunoreactivity differed across histological states in EV in a receptor-specific manner. The most consistent differences were lower TLR3 and TLR9 H-scores in cSCC than in non-neoplastic EV tissue and higher TLR9 H-scores in flat warts than in lesion-free skin. The TLR4 cSCC association was significant in the primary analysis but was not statistically significant after covariate adjustment; none of the TLR5 primary contrasts or tumour-differentiation analyses met the FDR threshold. The diagnostic and prognostic significance of these tissue patterns remains to be established, as diagnostic performance and clinical outcomes were not assessed in the present study. Independent studies are needed to determine their reproducibility and biological significance.

## Supporting information

Online Resource 2

Online Resource 1

## Statements and Declarations

### Ethics Approval

This study was performed in line with the principles of the Declaration of Helsinki. Approval was granted by the Institutional Review Board of Hospital das Clínicas, University of São Paulo (protocol 50871415.2.0000.0068; approval no. 1.401.205, 3 February 2016). The requirement for informed consent was waived because the study exclusively used archived, anonymized specimens.

### Author Contributions

Mirian Nacagami Sotto: Conceptualization; Methodology; Validation; Investigation; Resources; Writing – Review & Editing; Supervision; Project administration; Funding acquisition.

Cibele Conceição dos Apóstolos Pereira: Conceptualization; Methodology; Investigation; Data curation.

Luis Alberto Ribeiro Fróes Júnior: Methodology; Software; Formal analysis; Investigation; Data curation; Writing – Original Draft; Visualization.

Lana Luiza da Cruz Silva: Investigation.

Naiura Vieira Pereira: Investigation.

Maíra Garcia Saldanha: Investigation.

Walmar Roncalli Pereira de Oliveira: Investigation; Resources.

### Data Availability

The datasets generated and/or analysed during the current study are available from the corresponding author on reasonable request.

### Supplementary Information

Online Resource 1 Distribution of TLR3 (a), TLR4 (b), TLR5 (c) and TLR9 (d) H-scores across lesion-free skin, flat wart, cSCC in situ and invasive cSCC grades 1, 2 and 3. Each point represents one analytical specimen H-score; for TMA tumours represented by multiple eligible cores, the plotted value is the median core H-score. Boxes indicate the interquartile range, central lines the median and whiskers the most extreme values within 1.5 times the interquartile range. Separation of invasive grades shows the distributions underlying the prespecified G1–G3 differentiation analyses

Online Resource 2 Complete inferential results for the 12 prespecified primary contrasts, tumour-differentiation analyses and sensitivity analyses

