## Supplementary figures and images for "Toll-like receptor expression across histological states of Epidermodysplasia Verruciformis"

### Online Resource 1

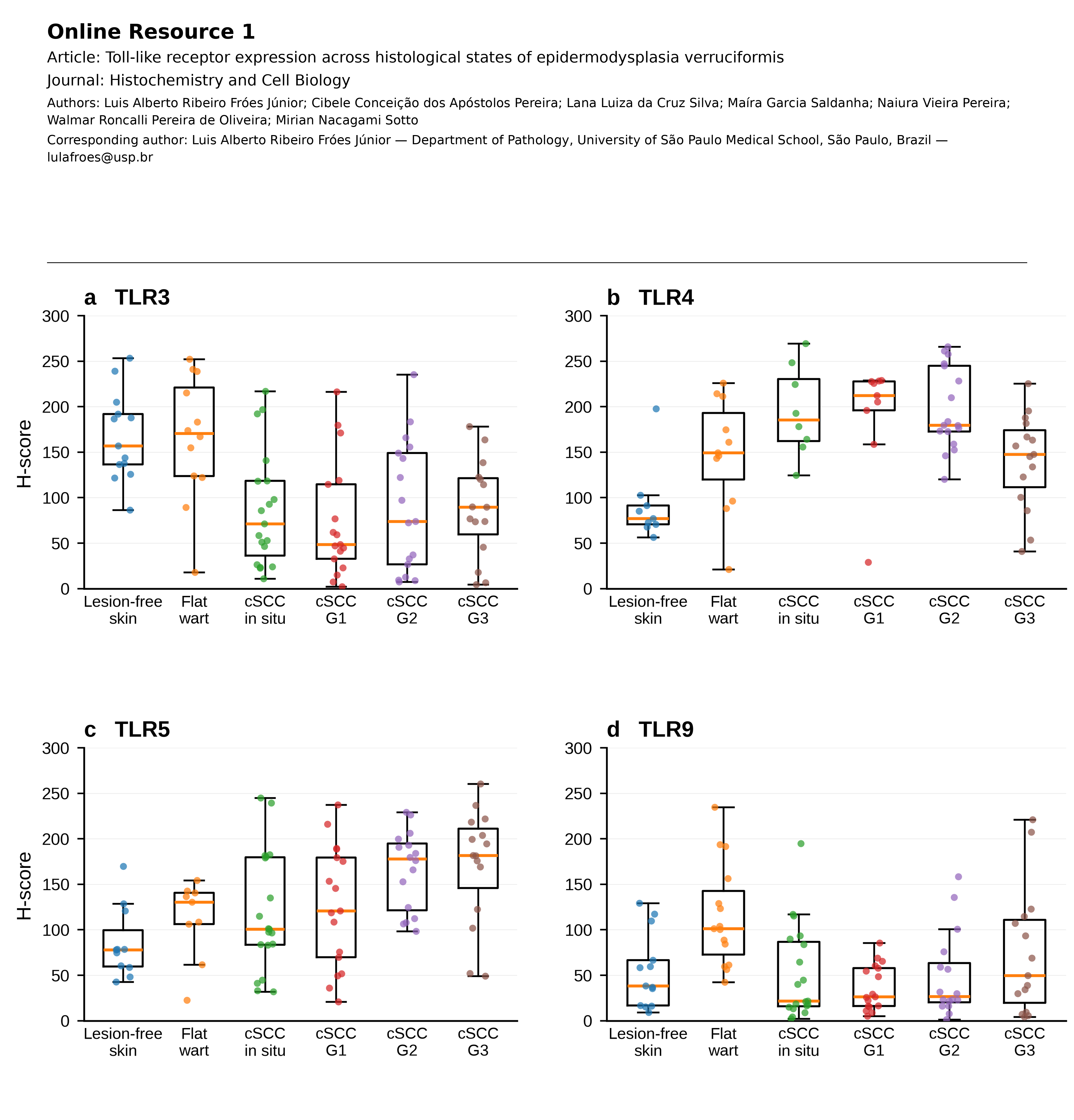
